# Potential Impact of Climate Change on One-Horned Rhinoceros (*Rhinoceros unicornis*) in Nepal

**DOI:** 10.1101/2020.05.04.076562

**Authors:** Ayush Adhikari, Deep Narayan Shah

## Abstract

Abrupt change in climate or simply termed as climate change is considered to be one of the major challenges in biodiversity. Change in climate has impacted many species around the world, particularly threatened species like One-Horned Rhinoceros (*Rhinoceros unicornis*). *Rhinoceros unicornis* is placed as an endangered species by International Union for Conservation of Nature (IUCN). Being an endangered species, studies regarding the impact of climate on the distribution of *Rhinoceros unicornis* is very rare in Nepal. Thus, the present study focuses on identifying the potential impact of climate change on the suitable habitat of *Rhinoceros unicornis* in Nepal using Species Distribution Modelling (SDM). For this, we used the present climatic scenarios and two greenhouse concentration trajectories (RCP 4.5 and RCP 8.5) for two different time periods (2050 and 2070) using different bioclimatic variables. Our model demonstrated the loose of the suitable habitat of *Rhincoeros unicornis* by 51.87% and 56.54% in RCP 4.5 for year 2050 and 2070 respectively. Under RCP 8.5 for year 2050 and 2070, the model demonstrated the loose of present suitable habitat by 54.25% and 49.51% respectively. Likewise, our result also predicted elevation as an important bioclimatic variable. This study would provide an information to the policy makers, conservationist and government officer of Nepal for the management and protection of habitat of *Rhinoceros unicornis* in present and future climatic context.

## Introduction

The climate of Earth is changing in an unprecedented way. The study of the Intergovernmental Panel on Climate Change (IPCC) suggests that within the time frame of 1880-2012, the global average temperature of Earth had increased by 0.85 °C (1). And, it is further predicted to increase the temperature by minimum 0.3°C-1.7°C under Representative Concentration Pathways (RCP) 2.6 and maximum by 2.6-4.8°C under RCP 8.5 scenario (1). Such change in climate has major challenges in biodiversity [(2),(3)]. Change in climate has widely affected the environment and its impact is seen worldwide, creating global signal of climate-induced range shifts and phenological responses crossing all the ecosystem and taxonomic groups [(4–8)]. The condition is likely to amplify in the future placing the terrestrial and aquatic biodiversity in great pressure (5).

Nepal lies in Himalayan region and due to its geographical isolation and species range limitations, the country has become one of the most vulnerable regions to climate change (9, 10). Glancing at the data of 25 years from 1982-2006, the temperature in the Himalayas has warmed up by around 1.5 °C, which is three times more than the global average temperature (10). Alarming warming in the Himalayas has started to exhibit in form of melting glaciers, change in hydrology patterns, agriculture, biodiversity, human health, ecosystem and livelihood (2). Such changes in climate have placed a threat for the large mammals of the mountainous country like Nepal (11, 12). Effects have started to seen through the fragmentations and reductions of the suitable habitats of the animals. Numerous studies such as (13–15) have predicted the increase in temperature and precipitation rate in the Himalayas. Such an increase in temperature might have a profound effect on biodiversity and ecosystem, for example, predicted for Marco Polo Sheep and Gaint Panda. It is predicted that Marco Polo Sheep would lose its suitable habitat area in lower elevation of Tajikistan (12). Same was the case for Red Panda, in which model predicted the loss of favorable habitat by 16.3 ± 1.4 (%) in China. However, the distribution of *Rhinoceros unicornis* for the present and future climate scenarios is still unknown. No research has been carried out to date for identifying the habitat change of *Rhinoceros unicornis* under the present and future climate change scenarios.

The *Rhinoceros unicornis* (*photo 1*), which is also termed as flagship and an umbrella species, if conserved and protected could support in the protection of other naturally co-occurring species (16, 17). Due to the unique habitat requirement of the Rhinoceros, they like to remain in the alluvial floodplains dominated by the sub-tropical climate vegetations and availability of water and grasses for year-round (18). Once found in the Indian sub-continent along the flood plain of Indus, Ganges, Sindh river from Myanmar in East and Pakistan in the west. Currently, *Rhinoceros unicornis* is limited in the small pocket area of South Asia region, especially in lowland national parks of Nepal and India. In the national context, there are around 645 individuals (19) in wild and are classified as endangered species of Nepal by the International Union for Conservation of Nature (IUCN). The inadequacies of the favorable habitat followed by anthropogenic activities are considered as one of the big challenges for the conservation of geographically restricted species like *Rhinoceros unicornis* (20). Furthermore, suitable habitats of the *Rhinoceros unicornis* are declining due to human disturbance and climate change (21). The climate change has always identified as a threat to global biodiversity especially for terrestrial species (22). Thus, a study on the distribution of the species and the identification of the climatically favorable habitat area are considered an essential component for long term conservation of species like *Rhinoceros unicornis*. The conservation action plan of *Rhinoceros unicornis* prepared by the Government of Nepal, highlighted climate change as a major threat to the survival of species (19). However, there is no research on identifying the impact of climate change on *Rhinoceros unicornis* so far. Majority of the researches have only been focused in human-wildlife conflict (23–25), demography (26) and effect of invasive species (27, 28). Limited studies have stepped up and focused on identifying the Population Viability Analysis of this species (20) and modelled its habitat distribution in Orang National Park (29). However, there is a paucity of information on identifying the potential impact of climate change on *Rhinoceros unicornis* and its habitat, especially in Nepal. In recent years, there is a trend of using various statistical modelling methods for predicting the potential distribution of species in time and space (30). Here, we modelled the current habitat distribution of *Rhinoceros unicornis* in Nepal and identified the potential habitat of species in changing climate for different representative concentrative pathways. Our work is the first kind of study in Nepal that has used ensemble modelling technique for predicting the habitat suitability of *Rhinoceros unicornis* in Nepal under climate change. We used different bio-climatic variables along with physiographic layers for understanding the distribution of species under climate change scenarios. Such findings would be a great source of reference for the conservation policies in Nepal.

**Photo 1:**
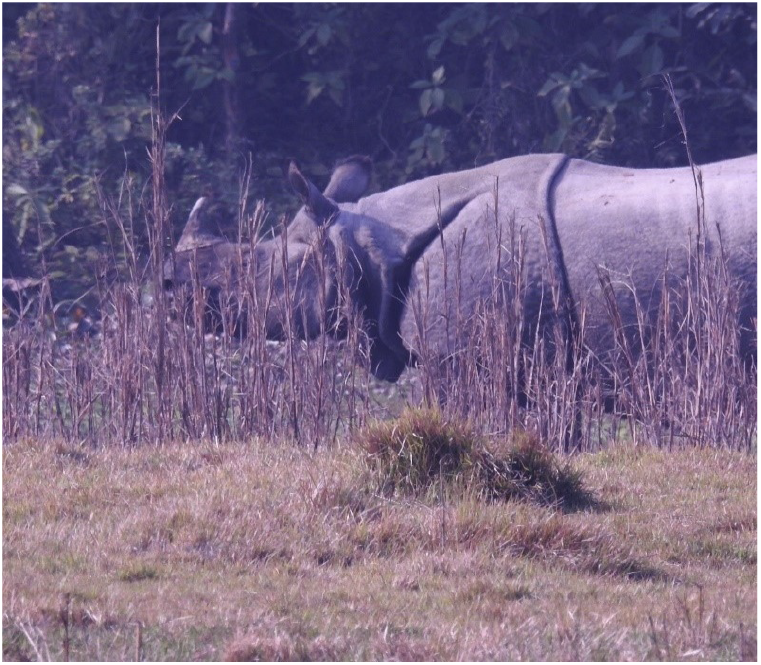
One Horned Rhinoceros (*Rhinoceros unicornis*)

## Methods

### Study Area and Presence Data

Nepal occupies an area of 1,47,181 km^2^ and stretches between 26.36° N–30.45° N and 80.06° E– 88.2° E within an elevation of 60-8848 m above sea level. The country is divided into five physiographic regions; Lowlands (Siwaliks and Terai), Hills, Mid-mountains and High Mountains. *Rhinoceros unicorns* are mostly found in the lowlands of Nepal particularly in the three protected areas; Chitwan National Park (CNP), Bardia National Park (BNP), Parsa National Park and Shuklapantha National Park (19).

The occurrence points of *Rhinoceros unicornis* was collated from the entire range i.e., lowland national parks of Nepal from different sources such as national census reports, scientific literature, personal communication and via geo-referencing of maps (28, 31). At first, we collected 487 numbers of occurrence points and laid down in the 1km*1km grid cells of Nepal. Then, we removed all the duplicate records and multiple sightings within the grid cells. The primary motive of removing duplicate and multiple records were to build spatially filtered data. Such spatially filtered data prevents over fitting of the result and helps in better model performance (32). Out of 487 occurrence points of the species, 372 presence points were selected for the modelling*s*. These occurrence points are not shown in map because *Rhinoceros unicornis* is protected species in Nepal so publishing occurrence points of this species might create conservation problems.

### Environmental Variables

For performing the modelling of the *Rhinoceros unicornis,* we used 19 bioclimatic variables obtained from worldclim data sets (www.worldclim.org) (33). The physiographic layer like elevation was also downloaded from the worldclim data set. The slope and aspect data set were further prepared from the elevation data set. For avoiding multicollinearity among the selected variables, we performed collinearity analysis. Based on the expert knowledge and through the support of literatures, among all 22 variables, each pair of highly correlated variables (r >0.7) was removed. The variables like altitude, mean annual temperature and precipitation are considered as important bioclimatic variables for *Rhinoceros unicornis* (34, 35). So, these above-mentioned variables along with other sets of bioclimatic variables like annual precipitation, temperature of annual range, annual mean temperature, isotheramality, precipitation of driest quarter, precipitation of seasonality, precipitation of driest month, elevation, slope and aspect were selected for modelling. The selected variables were under the resolution of 0.5 arc seconds (approximately 1 km^2^).

For the future distribution of species, we used the data from the Beijing Climate Center Climate System Model (BCCCSM1.*1*). We downloaded the Beijing Climate Centre Climate System Model data for RCP 4.5 and RCP 8.5 for two years 2050 and 2070. We choose BCC CSM 1.1 over other various models because of its accuracy in simulating the current climate of the Tibetan Plateau (36). Since Nepal is at proximity to Tibet so it makes sense to use output based on BCC CSM1.1. RCP 4.5 scenario is considered a stable scenario where it predicted to minimize the emission of greenhouse gases while the RCP 8.5 scenario is considered a worst-case scenario where greenhouse emission is expected to increase in the atmosphere.

### Species Distribution Modelling

Several modelling techniques are available for predicting the distribution of species in different series of time. Various algorithms available today are used for describing the species-environment relationship (37). Generally, the production of the habitat suitability map depends upon the algorithm used during the process of modelling (37). We used the BIOMOD2 package in R for performing the habitat distribution modelling of *Rhinoceros unicornis*. Unlike the single model algorithms, BIOMOD2 incorporates ensemble modelling algorithm and consider producing better accuracy (38, 39). We used six algorithms available within the BIOMOD2 package in R for building up the ensemble model. The used six algorithms were two regression methods (GAM; Generalized Adaptative Model, MARS; Multivariate Adaptive Regression Splines) and four machine learning methods (GBM; Generalized Boosting Model, ANN; Artificial Neural Network, RF; Random Forest and SRE; Surface Range Envelope). GAM uses a set of polynomials termed as smoothers for generating the curves by local fitting to the subsection of data (40). One of the distinct advantages of using GAM over the Generalized Linear Model (GLM) is that GAM can be used where the relationship between dependent variables is not linear (40). MARS is a regression model that works on the hypothesis that the model coefficients vary over different levels of explanatory variables (41). And it is a non-parametric procedure that draws a relationship between the dependent and independent variables (41). Similarly, GBM is a machine learning method and in the BIOMOD2 package this algorithm uses boosted regression trees (42). Under this algorithm, each of the individual models possesses regression trees. Likewise, ANN is also a machine learning method whose algorithm is handled by two basic parameters the amount of weight decay and the number of hidden units (43). Under ANN, BIOMOD2 packages use multi-layered feed forward neural networks, which are trained by back propagation algorithm (43). RF is based on the classification method and results are generated from classification trees (44). Data are classified into different classes based on homogeneity. BIOMOD2 package uses 500 trees and information is extracted from each selected variable (44). Finally, SRE is also a machine learning method that creates arrays of values for each environmental variable at the given presence points. It is the simple method for predicting the distribution of species and it directly predicts the presence-absences of species.

Like presence points of species, the absence points are also considered as valuable data in SDM. The absence data are considered as an important factor for SDM algorithms and model assessment techniques (45). Since, we did not have true absence points so we generated the pseudo absence point within the model by using SRE strategy. Under the SRE strategy absence points are selected in area by the models which are environmentally dissimilar from the presence points of species (46). Based on (47, 48) we used 10,000 pseudo absence points for the modeling of *Rhinoceros unicornis*. The pseudo-absence generation was repeated three times for preventing biasness in model. Similarly, we split our data into two different sets; 75% of the data were used for the calibration process and remaining 25% of data were used for testing data set.

There are not specific methods for evaluating the predictive performance of the model. Various literatures from (47, 49, 50)have used Receiver Operating Characteristic (ROC) and True Skill Statistics (TSS) for evaluation. Both, of the methods can be used independently but it is advisable to run them all for cross-comparisons (39). Here, we used TSS for evaluating the predictive performance of model while ROC and Kappa were used for cross-checking the predictive performance of model. TSS considers both omission and commission errors, and value ranges from −1 to +1. TSS value +1 indicates perfect model, −1 indicates failure of model while values from 0.7 to 0.9 is considered as good model (39, 51). We selected TSS score ≥ 0.7 for building up the ensemble model from the projection of six different algorithms by using weighted mean approach. Weighted mean approach builds up the model based on the selected threshold of TSS score and provides more robust predictions than other consensus methods (52). The output generated from weighted mean approach is presence absence map which was further classified into three different classes of habitat suitability; High Suitability (greater than 80% of probability), Medium (60-80% of occurrence) and Low (40-60% of occurrence). For, calculating the change in habitat of *Rhinoceros unicornis* we used biomod range size function in BIOMOD2 package.

## Results

### Model Performance and Variable Importance

Overall, TSS score of the ensemble model was 0.95, suggesting predictive distribution of *Rhinoceros unicornis* with high level of accuracy. The average TSS score GAM was 0.93, MARS with 0.91, GBM with 0.93, RF with 0.93, ANN with 0.89 and SRE with 0.72. Similarly, we used evaluation strip built within BIOMOD2 package for calculating important bioclimatic variables. Result on the bioclimatic variables showed elevation (19.79%) as important variable. Similarly, seasonal precipitation (18.94%), annual precipitation (15.03%), temperature of annual range (13.05%) and mean annual temperature (12.41%) were considered as top four important bioclimatic variables for predicting the habitat distribution of *Rhinoceros unicornis* in different series of time and scenarios. Likewise, other important bioclimatic variables were Aspect (1.39%), Isothermality (2.03%) and Slope (3.69%).

**Figure 1:**
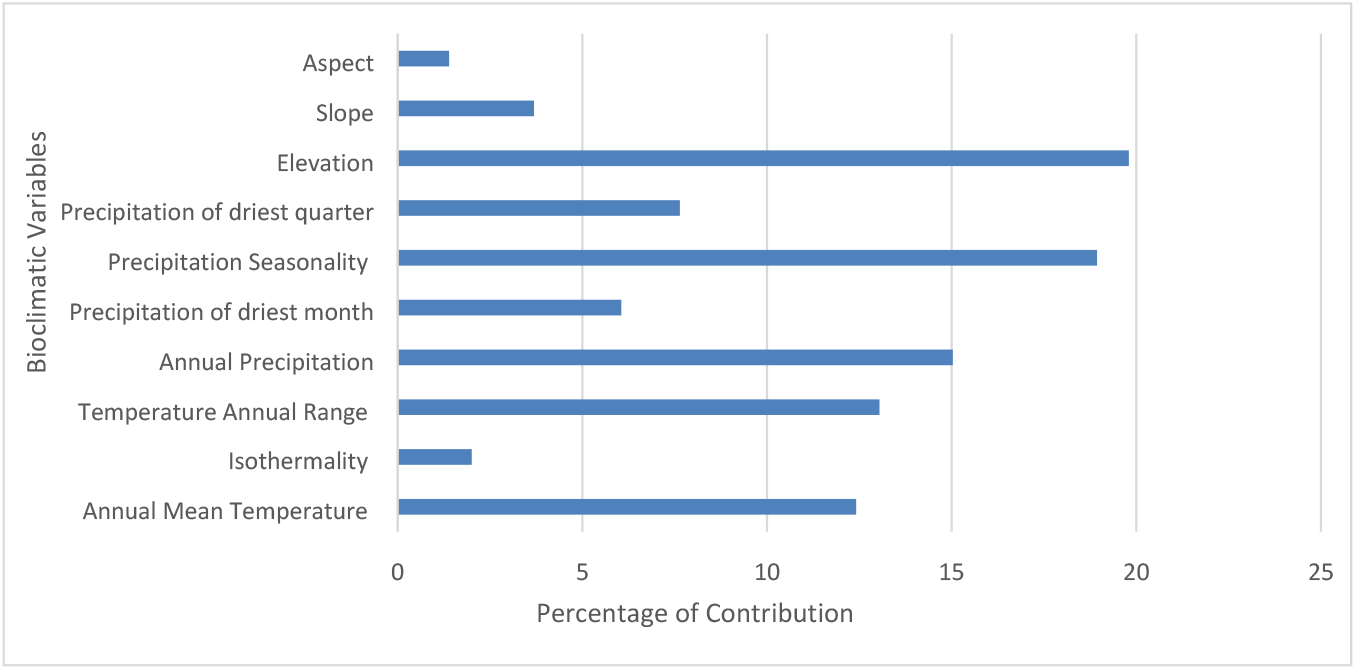
The important bio-climatic variables identified for predicting the distribution of suitable habitat of *Rhinoceros unicornis* in Nepal.

### Current Distribution of Rhinoceros unicornis

Our model estimated that *Rhinoceros unicornis* nearly occupies 7,240 sq. km of total area of country, which is approximately 5.01% of total land (fig. 2). Habitat of this species is mostly concentrated within the lowland national parks of Nepal from Shuklaphnata National Park to Chitwan National Park. Model predicted that, 659.09 sq. km of CNP is climatically favorable for the presence of *Rhinoceros unicornis* (Combing all the probabilities; high, medium, low). Similarly, 578.19 sq. km and 353. 69 sq.km of BNP and SWR is climatically favorable for the presence of *Rhinoceros unicornis*. While, remaining of the habitat suitable areas of *Rhinoceros unicornis* lies outside the national parks especially in buffer zones and community forests. Similarly, our model also predicted habitat suitability of this species in Eastern Nepal which is new identified area for *Rhinoceros unicornis*.

**Figure 2:**
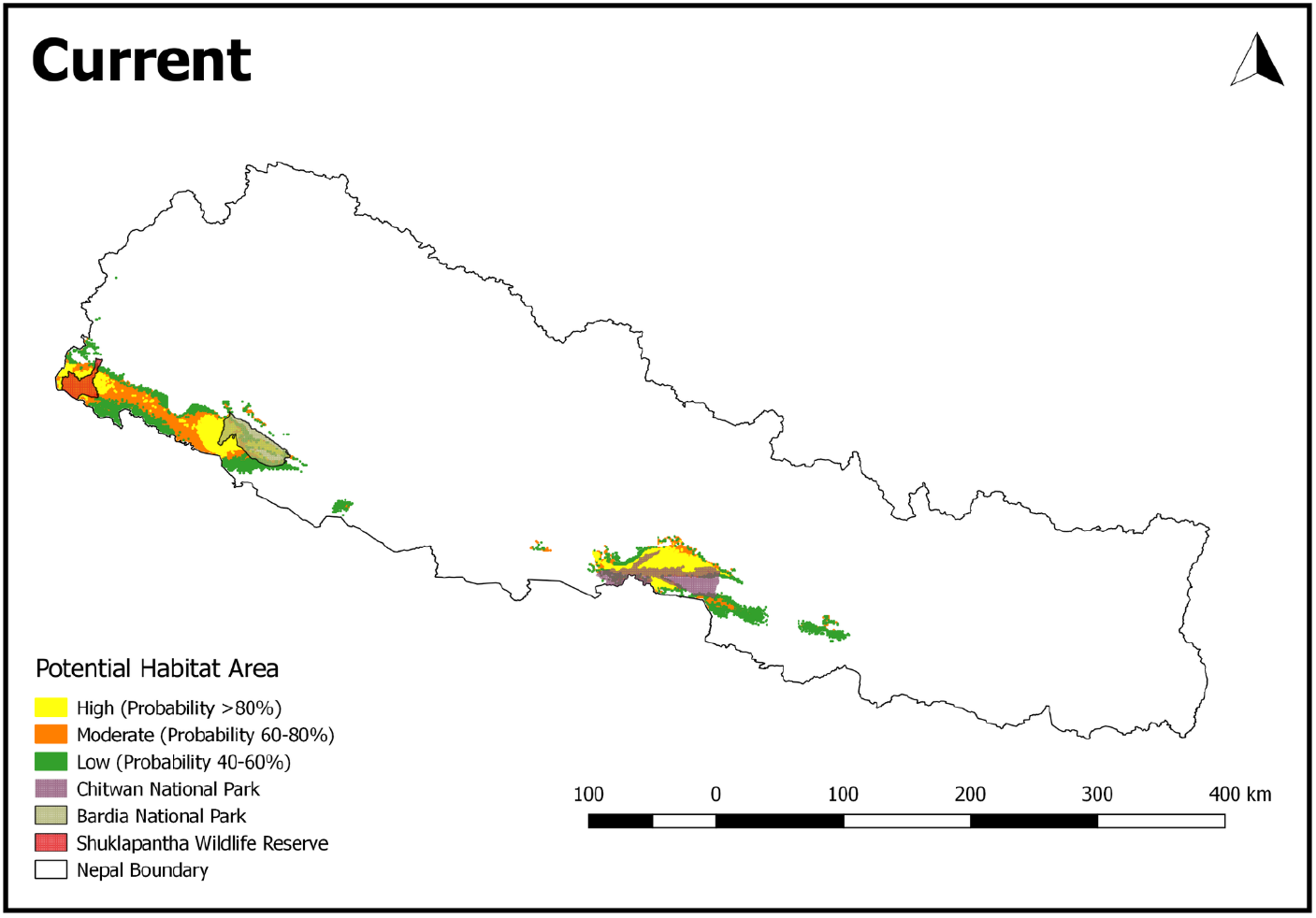
Map showing the Current Distribution of *Rhinoceros unicornis* in Nepal

For two different years under the two different climatic scenarios, *Rhinoceros unicornis* is predicted to lose its current suitable habitat (fig. 3 a,b,c,d). As shown in figure 3 (a,b) and table 1, under RCP 4.5, our model predicted that species would lose around 639 km^2^ and 727 km^2^ of present habitat for both years 2050 and 2070 respectively. While this species would likely gain 477 km^2^ and 491 km^2^ of habitats for two consecutive years 2050 and 2070. Overall, under this scenario, 2050 and 2070, our model predicted that the species habitat would decrease by 13.149% and 18.295 % respectively.

**Figure 3:**
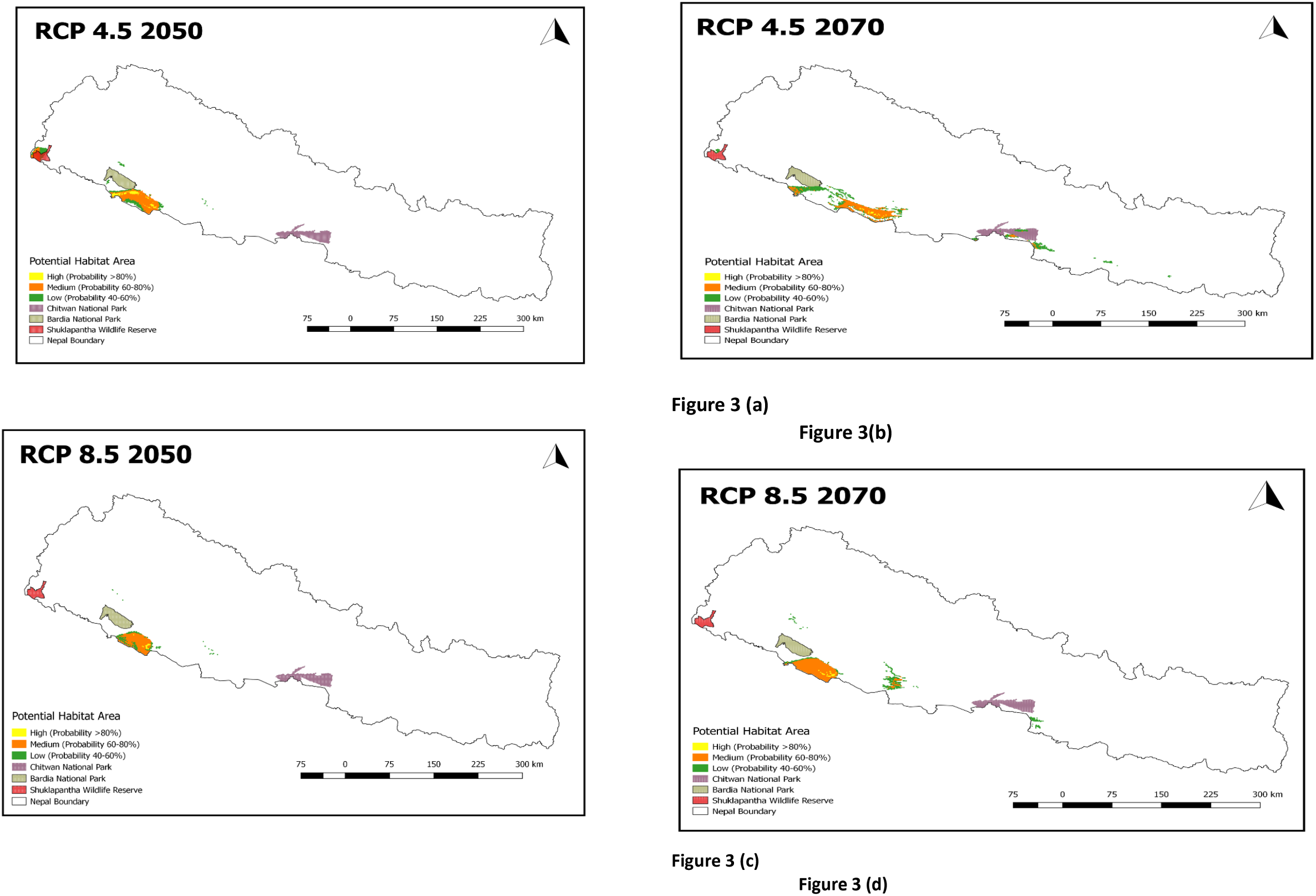
Change in potential suitable habitat area under different climatic scenarios of *Rhinoceros unicornis* in Nepal.

**Table 1:** Habitat Loss and Gain by *Rhinoceros unicornis* under different climatic scenarios

| RCP/years | Loss of Habitat | Gain of Habitat |
| --- | --- | --- |
| RCP4.5/2050 | 639 Km<sup>2</sup> | 477 Km<sup>2</sup> |
| RCP4.5/2070 | 727 Km<sup>2</sup> | 491 Km<sup>2</sup> |
| RCP8.5/2050 | 632 Km<sup>2</sup> | 455 Km<sup>2</sup> |
| RCP8.5/2070 | 606 Km<sup>2</sup> | 445 Km<sup>2</sup> |

Likewise, a similar result was obtained by the model under RCP 8.5 for both the years 2050 and 2070 Figure 3 (c,d) and table 1. The model predicted, *Rhinoceros unicornis* would lose its present suitable habitat area by 632 km^2^ and 606 Km^2^ in 2050 and 2070 respectively. For the year 2050, 54.23% of currently occupied sites would be lost whereas species is predicted to gain its new habitat by 39.06%. The scenario is the same for the year 2070, where the model predicted the loss of current habitat by 49.51% and gained new habitat by 36.356%. Overall, under RCP 8.5, our model has predicted the maximum reduction of species from central and in the far western part of Nepal especially in Chitwan National Park and Shuklaphanta National Park.

## Discussions

The species distribution modeling is considered a valuable tool for managing biodiversity, which includes an effort to conserve rare species, identify the biodiversity hotspots, biological response to climate change, predicting habitat suitability, finding out the problem of invasions and others (53). Such predictive modeling has become a valuable tool for conservation planning and the management of wildlife (54). Therefore, it is essential to understand the distribution of species in terms of ecological requirement and biological responses to present and upcoming climatic change through the use of species distribution modeling procedures (55). We modeled and predicted the distribution of *Rhinoceros unicornis* in Nepal under different climatic projections in different series of periods by including a set of bioclimatic variables. The ensemble modeling technique for performing species distribution modeling in the BIOMOD2 package is considered a powerful tool for predicting the potential habitat of species. However, such technique is also associated with the uncertainties. Here, we have reduced uncertainties in modeling technique through cross-validation procedure in which same modeling data sets were used to construct and evaluate the model. At first, 75% of model data sets were used in calibration and the model was evaluated by the remaining 25% of data sets using True Skill Statistics (TSS). Furthermore, the overall TSS score value was 0.95, which further justifies that predictive distribution of *Rhinoceros unicornis* is statistically correct.

The elevation is considered an important variable for the distribution of *Rhinoceros unicornis* in Nepal. This fact is also supported by (29) who considers elevation as a key factor for the distribution of species. Likewise, seasonal precipitation, annual precipitation, mean annual temperatures are considered as other important variables for the distribution of species (35, 56). In Nepal, *Rhinoceros unicornis* is mostly distributed in the lowland’s national parks of Nepal like Chitwan National Park, Bardia National Park, and Shuklapantha National Park. The simulated current projection of *Rhinoceros unicornis* is highly correlated with its actual distribution, which has predicted the distribution of species within the lowland of Nepal. Previous research from (25, 28) had recorded species from Kanchanpur in Far-western Nepal to Chitwan in central region Nepal within an altitude of 156 m to 820 m. Our model also identified these areas as a climatically suitable area for the distribution of species. Within this range, *Rhinoceros unicornis* occupies an area of 7,240 km^2^, which is 5.01% of the total area of the country. Similarly, our model predicted the distribution of species from the buffer zone of CNP where there is the problem of human-rhino conflict (57, 58). In addition to this, the model also predicted the distribution of species in the Eastern region of Nepal (Rautahat and Sarlahi). This area is hypothesized as a suitable habitat area by Rimal *et al* (34).and the dispersal of species had occurred in recent years in these areas. Likewise, our model also predicted lower elevation of the mid and far western part of Nepal as a suitable area for the distribution of species. Kailali and Banke’s region is considered a favorable area for the distribution of species. In the past, these areas were climatically favorable for *Rhinoceros unicornis* as their distribution was widely distributed in the floodplains of the Ganges and Brahmaputra Rivers of India (59). But, in the present context due to the human interference and construction of physical infrastructures like road, settlement have hindered the distribution of species within this range. Nevertheless, if a study regarding the conservation of the habitat of *Rhinoceros unicornis* is carried out in future, this area would serve as an excellent candidate for the inspection.

Similarly, our model predicted the loss of present suitable habitat of *Rhinoceros unicornis* in different representative concentrative pathways 4.5 and 8.5 under two different years 2050 and 2070. The majority of the losses were observed from central Nepal especially from the Chitwan National Park where the occurrence points of the species were mostly concentrated. Such a decrease in the favorable habitat of the *Rhinoceros unicornis* might be due to the selection of environmental variables. Our model suggested precipitation of seasonality and mean annual temperature as important bioclimatic variables. Temperature and precipitation have always been crucial factors for the distribution of *Rhinoceros unicornis*. Numbers of sophisticated climate models have predicted the rise and fall in precipitation and temperature level within the end of this century (60). By the end of the 2090s, average annual temperature in the Himalayas might raise by 3.0-6.3°C (61). It is predicted that within an average raise of temperature by 2.3 °C, the amount of precipitation rate would be increased by 5.2%(62). In case of Nepal, an increase in temperature would increase in the rate of perceptible water by 20-24% (63). Similarly, monsoon precipitation is expected to increase by 4-12% in short term and 4-25% in long term (64). Such increases in precipitation rate would increase the river runoff by approximately 7.3% creating major flood events throughout the Indian Subcontinent (62). An increase in flood events in the future within the lowland of Nepal might create the problem of soil erosion, the cutoff of topsoil and sediment deposition within the grassland of Terai. Such deposition of sand content and cutoff of the top soils might reduce the productivity of riverine ecosystem where *Rhinoceros unicornis* inhabits. A recent example was the flooding event that took place in Chitwan National Park and Kaziranga National Park, which caused the mortality of *Rhinoceros unicornis* and even caused the dispersion of this species from habitat. This scenario could be exaggerated in the future causing the dispersion of *Rhinoceros unicornis* from its current favorable habitat. Unlike the precipitation, warming is projected to increase by 2.2–3.3 °C for RCP4.5 and 4.2–6.5 °C for RCP8.5 (64). Such increase in temperature might dry up the water resources in low land of Nepal causing the habitat shift of *Rhinoceros unicornis*. This line of fact is also supported by (65) who came up with conclusion that under the condition of low availability of water or drought most of the ungulates in Africa either shifts their range or faces extinction. *Rhinoceros unicornis* have direct relationship with the stream sources and water availability. Generally, in national parks this species is particularly found within water containing areas like rivers, ponds and lakes (28, 66–68). The future increase in annual range temperature in Nepal might dry up the water resources and this could be one of the reasons for dispersal of this species from present habitat. Lower mid-west part of Nepal could hold good proportion of water in future due to the favorable temperature and in search of favorable habitat *Rhinoceros unicornis* might disperse to these areas.

Similarly, another possible reason for change in favorable habitat of the species might be due to the problem of invasive species. At the present scenario, invasive species like *Mikania micrantha* inhabits in most of the lowland national parks of Nepal especially in the riverine forest and alluvial grassland (69, 70). Study suggests that *Mikania micrantha* is spreading in aggressive way and has created problem in component of biodiversity like in forest cover, water availability, agriculture and even in grassland (66, 69, 70). Rise in temperature supports in dispersion of the invasive species to the greater range (71). In high gas emission scenario, invasive species tends to occupy or increase their habitat range by two-fold by 2070 (71). Our model has exhibited annual mean temperature and annual range temperature as important bioclimatic variables. Rise in temperature by 2-3° C and increase in the CO_2_ level by 600 ppm in RCP 4.5 and 1250 ppm in RCP 8.5 for both years (2050 and 2070) might make environment favorable for this invasive species and help in expanding its range. The expansion of this invasive species might spoil the grassland, shrubland and water quality of the region where *Rhinoceros unicornis* lives and even might reduce the carrying capacity of park (70). This might be another possible reason that might cause range contraction of this species in future which drives species to new favorable habitat.

Our model predicted the climatically suitable habitat of *Rhinoceros unicornis* in current and future climate only by using abiotic factors. Use of only abiotic factors for predicting the distribution of species have been criticized by (72) because there are others climate related stresses that impact in the distribution of species. Using only abiotic factors like temperature and precipitation would not be sufficient for projecting distribution of species in different series of time. However, using abiotic factors in predicting the habitat distribution of species is considered as crucial approach(73) for understanding the effects of climate change. Though, we have used different bioclimatic variables for predicting distribution of species, incorporating variables like distance to road, distance to water bodies, landcover, vegetation, soil moisture might add additional value to our research. Moving beyond this constraints, future studies of the *Rhinoceros unicornis* need to incorporate inter and intra species interactions among the species in the model.

## Conclusion

The study highlights the potential impact of climate change on distribution of *Rhinoceros unicornis* in different time scale under the present and future climatic conditions. There remains huge possibility of losing favorable habitat by the species on different gas emission scenarios by shifting its range to mid-west part of Nepal. Similarly, model also exhibits new areas of Nepal like Rautahat in East and Banke in West of Nepal as climatically suitable area for *Rhinoceros unicornis,* which could be the priority area for detail study for planners and conservation officers. In future climatic scenarios, CNP and SWR is expected to lose favorable habitats of *Rhinoceros unicornis*. Overall, the result derived from this research could act as baseline for the conservationist and policy makers for the conservation and habitat management of *Rhinoceros unicornis* in present and future.

